# MeNu GUIDE - a metabolite nutrition graph to uncover interactions with disease etiology

**DOI:** 10.1101/2024.10.12.618040

**Authors:** Vivian Würf, Josch K. Pauling

## Abstract

The relationship between diet and disease is well-documented, yet the complex interactions among foods, metabolites, and genetics makes research challenging. This study explores the potential insights offered by a knowledge graph that connects nutrition and diseases on a metabolic level. Ten ontologies and data from six databases were merged, resulting in a graph with over 25 million triple statements, stored in a Turtle file and added to a GraphDB repository. SPARQL queries revealed biases towards specific foods and conditions within the integrated databases. Despite these biases, this knowledge graph serves as a proof-of-concept, demonstrating the feasibility of integrating information from diverse resources to yield valuable insights and enabling the drawing of meaningful conclusions. The graph allows efficient identification of disease-related compounds and their food sources and enables the exploration of changes in metabolite concentrations, such as those occurring during food processing. Researchers could use such a knowledge graph to identify biomarkers, help generate new hypotheses, and improve experimental designs. Expanding the graph with automated text-mining and recipe data would further enhance its utility for nutrition research. Such a resource could advance understanding of the molecular mechanisms behind diet-disease relationships, guiding more targeted interventions.

## 1 Introduction

Nutrition research grapples with multifaceted challenges that render it inherently complex. It is well-established that many diseases have a strong connection to nutrition, including obesity [1], cardiovascular conditions [2], type 2 diabetes mellitus [3] and non-alcoholic fatty liver disease [4]. This highlights the vast scope of how dietary patterns influence health and disease. Certain diets can directly contribute to disease development, such as alcohol consumption leading to alcoholic fatty liver disease [5], while others can be used therapeutically, like exclusive enteral nutrition in the treatment of Crohn’s disease [6, 7]. However, the diversity of dietary patterns in different populations makes it challenging to generalize findings across varying cultural, socioeconomic, and geographical contexts. Additionally, conducting controlled experiments in nutrition research is challenging due to the long-term nature of diet-disease relationships.

Despite significant advancements, much remains unknown, especially regarding diet-related disorders and diseases. Conditions like Irritable Bowel Syndrome (IBS) and Inflammatory Bowel Disease (IBD) have reached a concerning level of global prevalence affecting millions of people worldwide [8–13]. However, our understanding of their underlying causes and optimal dietary interventions remains limited. This results in untapped potential for treatment, personalized interventions, supportive therapies, and disease prevention. While there is growing recognition of the role of diet in managing these conditions, tailored dietary strategies that effectively alleviate symptoms and improve the quality of life for patients are still lacking [14–17]. Despite considerable progress, the complexity of human metabolism and the multifactorial nature of disease etiology pose significant challenges to understanding the intricate connections between foods, metabolites, and diseases. Bridging this knowledge gap necessitates adopting innovative approaches to integrate information from heterogeneous data sources.

Multiple initiatives have been launched to compile comprehensive databases of food metabolites to understand their potential roles in human health and disease [18–20]. While the existing databases provide valuable information on the chemical composition of foods and their metabolites, gaps and inconsistencies persist. Additionally, they lack direct links to disease associations. Moreover, not every database utilizes standardized nomenclature and annotation for metabolites, which further complicates integrating data across different sources. As a result, while these databases serve as valuable resources for nutritional research, their limitations underscore the need for continued efforts to improve data quality, coverage, and interoperability to facilitate meaningful insights into the impact of food metabolites on human health.

Knowledge graphs are structured representations of data that capture and store entities and the relationships between them. They have emerged fairly recently as a promising tool for tackling complex biomedical problems. By organizing vast amounts of heterogeneous information from diverse sources such as scientific literature, databases, and electronic health records, knowledge graphs can provide a holistic view of biological systems. Researchers can then use graph analytics to extract meaningful insights and elucidate the intricate relationships between genes, diseases, drugs, and other biomedical entities. The success of knowledge graphs in biomedical research is evidenced by their diverse applications in areas such as drug repurposing, precision medicine, and clinical decision support [21–24]. Leveraging the wealth of interconnected data encapsulated within knowledge graphs enables researchers to navigate the complexities of biological systems, accelerate scientific discovery, and, ultimately, advance human health.

Here, we present the nutrition-disease knowledge graph MeNu GUIDE, which was constructed through a multifaceted approach that harnesses structured data from databases and ontologies (figure 1). Relevant data from various curated databases containing information on nutrition, metabolites, diseases, genes, and pathways were extracted and integrated to populate the knowledge graph. Our nutrition-disease knowledge graph offers a comprehensive and dynamic repository of interconnected information, facilitating easy, fast, and holistic exploration and analysis of the complex interrelationships between diet, metabolism, and health outcomes. By employing graph analytics, its utility extends beyond mere data aggregation, serving as a powerful tool for hypothesis generation and data-driven discovery. The ultimate goal of the nutrition-disease knowledge graph is to unravel the underlying mechanisms linking dietary factors to disease risk and progression, thereby informing evidence-based dietary recommendations and ultimately improving patient health outcomes.

**Figure 1:**
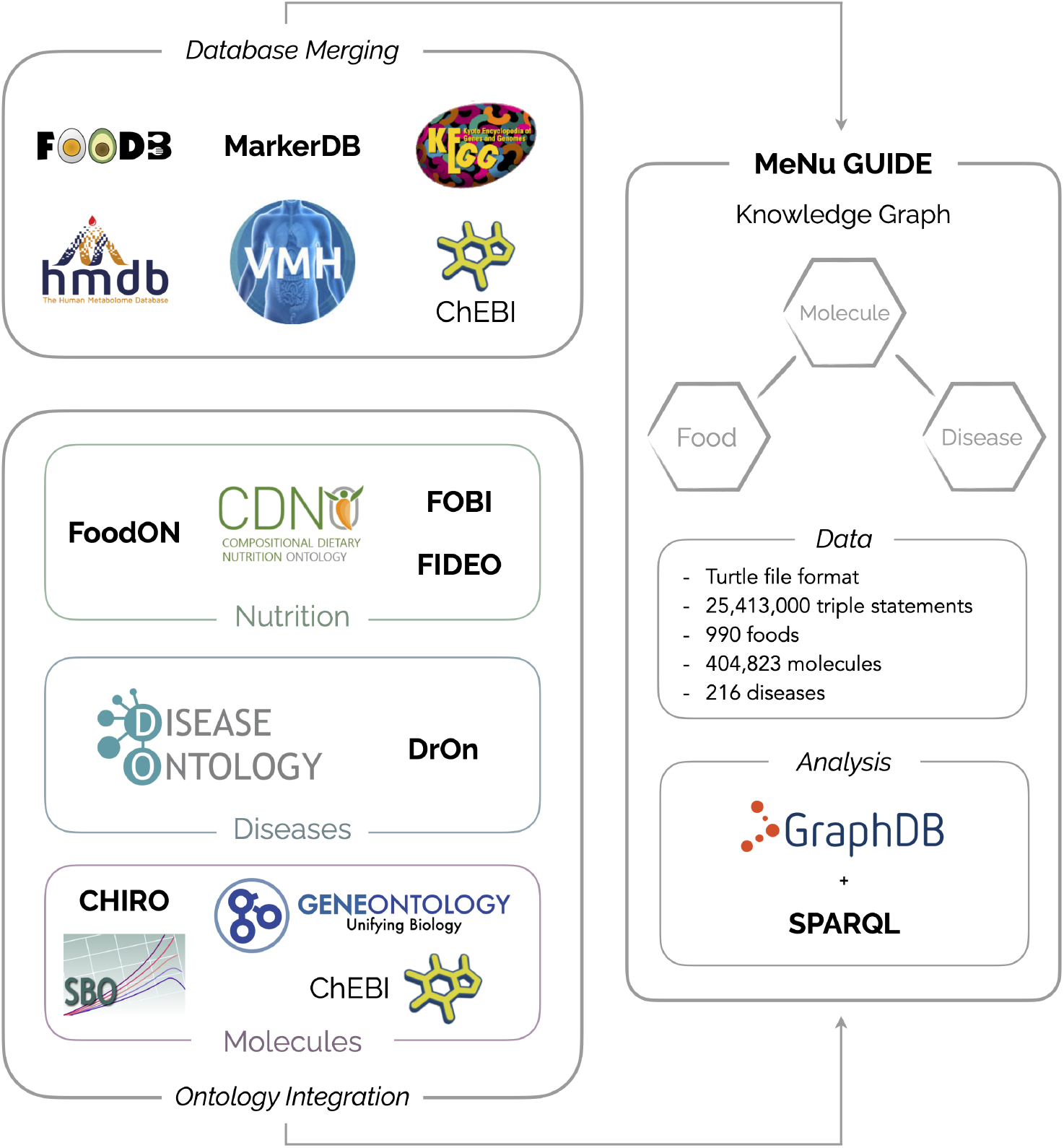
MeNu GUIDE is a knowledge graph containing over 25 million triple statements on the relationships between foods, metabolites and diseases. It was created by combining the information from databases with the knowledge stored in ontologies. MeNu GUIDE is available in the Turtle file format, which can easily be queried using SPARQL.

## 2 Methods

### 2.1 Database Merging

Information from six different databases was extracted and merged. The utilized databases include the Food Database (FooDB) [18], the Human Metabolome Database (HMDB) [25], MarkerDB [26], Chemical Entities of Biological Interest (ChEBI) [27], Kyoto Encyclopedia of Genes and Genomes (KEGG) [28] and the Virtual Metabolic Human (VMH) database [29]. Three of these databases (FooDB, HMDB and MarkerDB) are curated by the Wishart reasearch group at the University of Alberta in Canada and therefore form an ideal starting point for the database integration process. Beginning with **FooDB**, all relevant data was downloaded as CSV files from the website. Compounds in the Compounds.csv file were matched with their external descriptors in the CompoundExternalDescriptor.csv file to obtain as many identifiers as possible for merging with the other databases. In the next step, the hmdb_metabolites.xml file was downloaded from the **HMDB** website. Relevant information and identifiers were extracted for each compound and saved as a CSV file. The extracted HMDB compounds were then matched with the FooDB compounds based on the lowercase compound names, as it yielded the best results. To extract all compounds present in **MarkerDB** the file all_chemicals.tsv was downloaded from the website. All unique compounds were extracted and the API was queried to retrieve information on each compound. As each entry was associated with an HMDB ID, this information was used to match them with the previously merged FooDB and HMDB compounds.

To integrate the **ChEBI** database, the files containing information on compounds were downloaded from the website. Initially, only the highest confidence (3-star) entries were considered. However, references to 2-star entries in other databases necessitated the inclusion of all possible ChEBI compounds. Information about KEGG and International Chemical Identifier (InChI) identifiers were contained in separate files (’chebiId_inchi.tsv’ and ‘database_accession.tsv’), which were merged with the ‘compounds.tsv’ file. Discrepancies between ChEBI IDs in the HMDB XML file required manual verification and correction of duplicated IDs before merging the datasets. To extract all **KEGG** compounds, the provided REST API was utilized. Extracted flat files for each metabolite were parsed using regular expression, and KEGG compounds were matched to the dataframe via KEGG IDs provided by HMDB, FooDB, and ChEBI. **VMH** metabolites were extracted using the provided API. VMH compounds were matched using available VMH IDs, as well as HMDB, FooDB, ChEBI identifiers, and compound names. This resulted in the final table comprising all compounds present in the mentioned databases.

### 2.2 Ontology Integration

Ontologies were utilized as a systematic framework for organizing and harmonizing the diverse datasets extracted from various databases. For constructing this knowledge graph, ontologies describing nutrition, diseases, and compounds were integrated using the RDFLib library [30]. The nutrition-relevant ontologies included the Food Ontology (FoodOn) [31], Compositional Dietary Nutrition Ontology (CDNO) [32], Food Interactions with Drugs Evidence Ontology (FIDEO) [33] and Food-Biomarker Ontology (FOBI) [34]. For diseases, Human Disease Ontology (DO) [35] and The Drug Ontology (DrOn) [36] were integrated, and for molecules, ChEBI, ChEBI Integrated Role Ontology (CHIRO) [37], Gene Ontology (GO) [38, 39] and Systems Biology Ontology (SBO) [40] were included. Some ontologies already incorporate others, such as DO including ChEBI, which facilitated the integration process. For ontology alignment, precomposed and curated mappings were extracted from BioPortal [41] using its API. The final merged ontologies were saved as a Turtle file.

### 2.3 Combining Database and Ontology Information

The data composed from the database extraction was then added to the merged ontologies. To match the compounds in the database to the ontologies, unique Internationalized Resource Identifiers (IRIs) were created for each molecule using a generated compound ID. The compound name was added as label and the available properties such as monoisotopic mass, chemical formula, InChI and SMILES were added using the already available predicates from the ChEBI ontology. For a detailed list of the utilized predicates see table S9. Wherever possible, the compound was connected to corresponding classes in the ChEBI ontology.

The 990 foods and dishes present in the downloaded FooDB data were first matched onto the FoodOn ontology using the common and scientific names, if available. This left 250 entities that could not be matched automatically, but had to be mapped manually. All properties used to connect the food nodes are described in table S10. The next step was to create the relation between foods and metabolites through the content entries provided by FooDB. For this purpose only the content entries were selected, that were associated with a source, unit and with a content greater than zero. The content entries were added as a separate node and then connected to the corresponding food and metabolites via their respective IRIs. All properties associated with the content nodes are shown in table S11.

Similarly, the data on biomarkers from MarkerDB was added in the form of separate measurement nodes to serve as the connection between compounds and diseases. To accomplish this, the conditions from the MarkerDB entries first had to be mapped to the DO ontology. A first automated attempt matched 143 diseases, leaving 150 conditions to be matched manually. A comprehensive overview over the relationships connected to the measurement nodes can be found in table S15.

Finally, a new node was created for each reaction. The substrates and products of a reaction were then linked via their MeNu GUIDE compound IRI. Nodes were also introduced for the corresponding enzymes and genes and connected to the reactions they are involved in. A complete list of all implemented relations can be found in tables S12, S13 and S14 for reactions, enzymes and genes respectively.

The final Turtle file containing all triples was then loaded into GraphDB Version 10.6 [42] to create the final knowledge graph. The content of the graph was then explored using SPARQL Protocol and RDF Query Language (SPARQL) queries.

## 3 Results and Discussion

### 3.1 Evaluation of the connection between foods and compounds in the MeNu GUIDE knowledge graph

The final graph consists of 25,413,000 triple statements and contains information on 990 foods, 404,823 compounds, 216 conditions, 223,540 content entries and 2,987 biomarker measurements. To obtain an overview over the relations between the graph contents, highly connected nodes were extracted for each node type. Starting with the food nodes, query 1 was used to count the number of associated content entries. These content entries connect foods and compounds and originate from the FooDB database. One example of such a content entry and its connected nodes is visualized in figure 2. Each of these content entries is always associated with a food and compound. It’s properties include the content amount and corresponding unit. Additionally, it also mentions the reference, e.g. database or publication, from which this content information derives. For example, for content_1 the reference is “DTU”, which corresponds to the Danish food database Frida [43] provided by the Technical University of Denmark (DTU). If available, the content node also includes a minimum and maximum amount measured. The size of a node in the figure depicts its Resource Description Framework (RDF) rank in the graph, which is a measure of interconnectedness.

**Figure 2:**
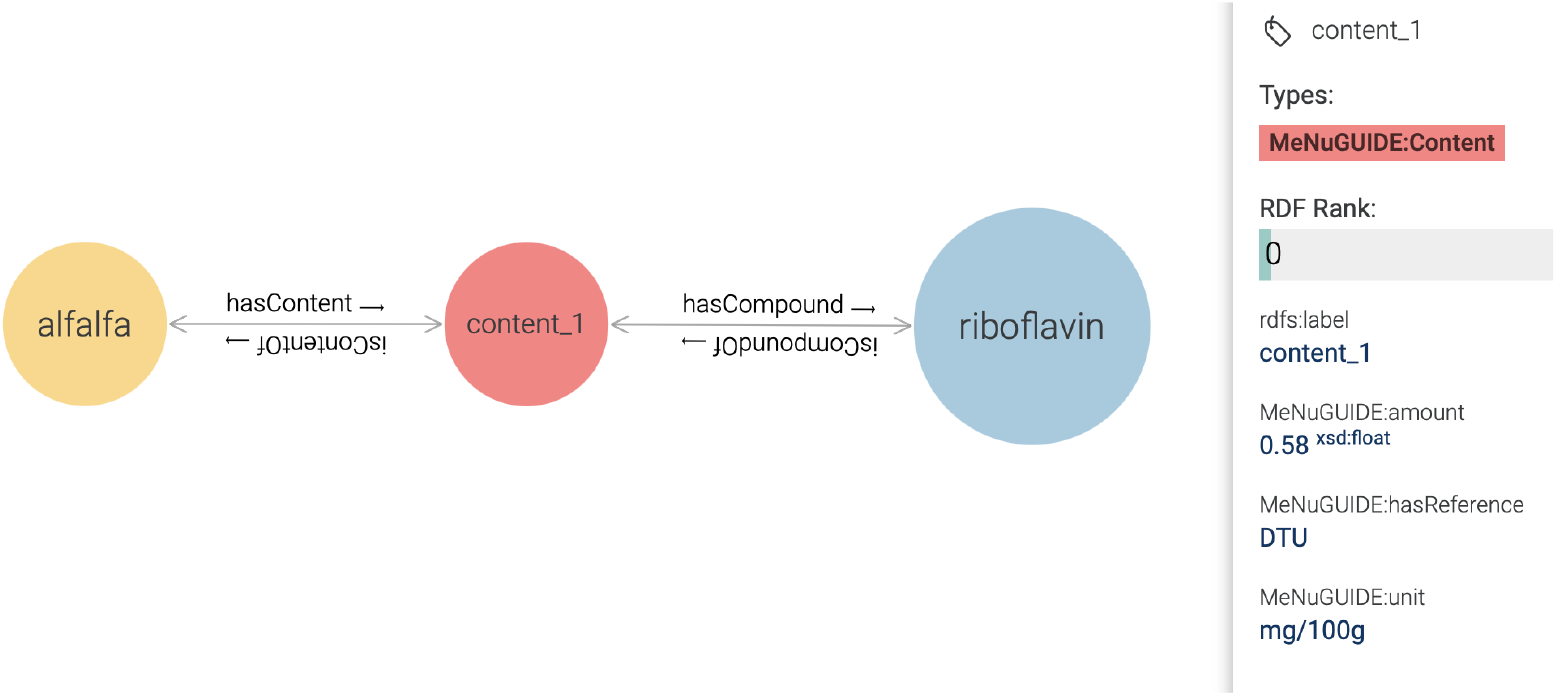
Example of a content node in the MeNu GUIDE knowledge graph connecting foods with the compounds they contain.

Table 1 displays the top 15 food nodes with the highest number of associated content entries. The top three most highly connected food nodes are cattle, cow milk and pork. The “cattle (beef, veal)” node is connected to 15,473 content nodes. However, querying for the compounds associated with these content nodes reveals that multiple content entries are connected to the same compound. This results in only 92 unique compounds associated with the “cattle (beef, veal)” node via the 15,473 content entries. Modifying the query to extract the number of distinct compounds associated with a food leads to major changes in the top 15 spots, except for milk, potato and corn, which also appear in table 2.

**Table 1:** Foods with highest associated content count.

| Nr | Food | Content Count |
| --- | --- | --- |
| 1 | cattle (beef, veal) | 15473 |
| 2 | milk (cow) | 10667 |
| 3 | domestic pig | 10310 |
| 4 | breakfast cereal | 6422 |
| 5 | baby food | 5002 |
| 6 | cheese | 4685 |
| 7 | soup | 3888 |
| 8 | sausage | 3413 |
| 9 | pastry | 2942 |
| 10 | biscuit | 2917 |
| 11 | common bean | 2599 |
| 12 | potato | 2527 |
| 13 | pizza | 2209 |
| 14 | corn | 2158 |
| 15 | snack bar | 1750 |

**Table 2:** Foods with highest associated compound count.

| Nr | Food | Compound Count |
| --- | --- | --- |
| 1 | milk (cow) | 1648 |
| 2 | carrot | 231 |
| 3 | corn | 223 |
| 4 | sweet orange | 186 |
| 5 | green bell pepper | 176 |
| 6 | beer | 175 |
| 7 | red bell pepper | 163 |
| 8 | potato | 161 |
| 9 | apple | 160 |
| 10 | broccoli | 154 |
| 11 | tea | 152 |
| 12 | black tea | 152 |
| 13 | green tea | 152 |
| 14 | parsley | 151 |
| 15 | mandarin orange | 148 |

In general, the highly connected food nodes are indicative for important agricultural industries. It is reasonable to assume that this is due to a research focus on foods in the most influential sectors. This is supported by looking at the Annual Survey of Manufactures compiled by the Bureau of the Census of the U.S. Department of Commerce. Data collected in 2021 shows meat processing to be the largest industry group in food and beverage manufacturing with 26.2% of sales, followed by dairy with 12.8% [44].

Similar trends are visible in the European Union, where about 39.8% of the total agricultural output in 2023 came from animals and animal products, mainly from milk and pigs [45]. This is estimated to value around C213.8 billion, representing an enormous market. Naturally, the contributions differ significantly between countries. In the case of France, cereals make up the largest share of the countries agricultural output in 2023 with 15.1%, shortly followed by wine with 14.3%. On the third and fourth spot are milk and cattle with 13.8% and 10.6% [46]. For Germany, milk made up the largest share of the agricultural output with 21.1%, followed by pigs with 12.6% [47]. Cereal is in the fourth spot with 11.9% and wine only makes up 2% of Germany’s agricultural output. Potatoes are listed separately from other vegetables in the country specific fact sheets and make up 6.0% and 7.2% of the France’s and Germany’s agricultural output. All this matches well with the highly connected food nodes and likely explains why they are so well researched.

Another reason for the overrepresentation of milk in the knowledge graph could be that the Wishart research group, which curates the FooDB and HMDB databases, also hosts the Milk Composition Database (MCDB) [48]. It is reasonable to assume that the MCDB shares data with the other databases in the ecosystem, which might lead to an unintended information focus on milk in this knowledge graph. This suggests that the knowledge graph must be enriched with information from additional sources to mitigate biases introduced during the database selection process. These sources could include more databases as well as the extraction of information from publications. Although this may not completely eliminate the research focus bias, it will help alleviate curation bias.

One advantage of the graph structure is that it allows for easy extraction of compounds that two or more foods have in common. To further illustrate this point, query 3 was used to compare the average content of compounds contained in milk and cheese, which is shown in figure 3. Four observations can be made based on this comparison which validate the knowledge graph content. One aspect is the ten-fold increase in sodium content of cheese which is likely due to the salting during the cheesemaking process. This helps to preserve the cheese and prevent the growth of pathogens [49]. Additionally, there is more lactose in milk compared to cheese. This is to be expected and consistent with the manufacturing process as only minimal amounts of lactose are left in the cheese after the metabolization to lactic acid by the starter culture during curd formation and removal of the whey, which retains about 90% of the lactose present [49]. Another observation is that cheese contains a lot more cholesterol than milk, in this case around 17 times more on average. The reason for this is likely the concentration process of the milk solids during cheese-making by removing water. As so much milk fat is retained in the cheese it is logical that this would also lead to an accumulation of fat-soluble vitamins present in milk. This fits with the increase in retinol and beta-carotene in figure 3. However, it is important to note that the levels of beta-carotene and other carotenoids, such as bixin and norbixin, may be elevated in cheese due to their possible use as food colorants [50]. The increase in compounds such as riboflavin, folate and vitamin B12 could be attributed to production by bacteria during the ripening process depending on which strains were used [51].

**Figure 3:**
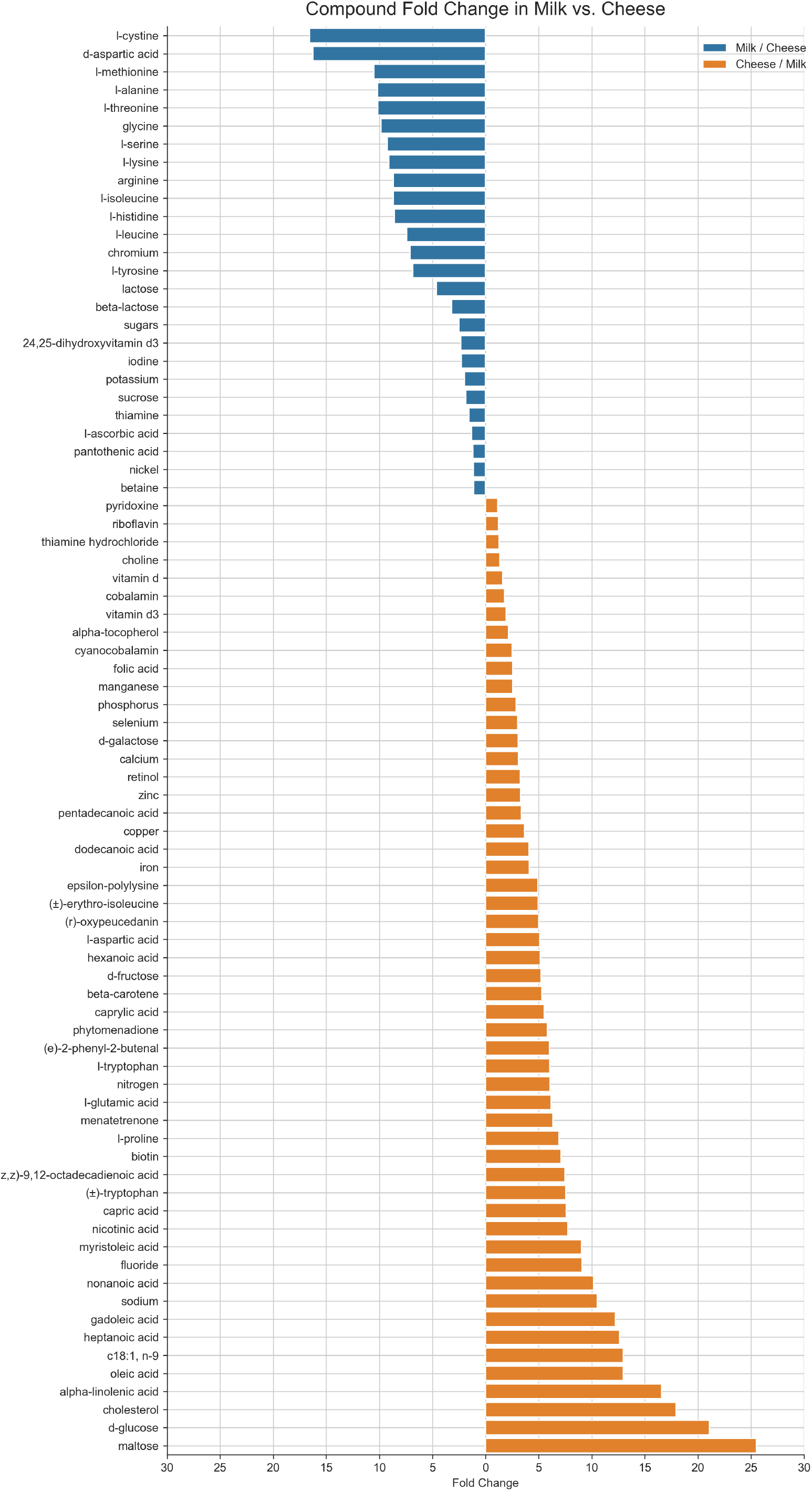
Fold change of compounds contained in milk and cheese found in MeNu GUIDE knowledge graph.

To further explore the graph contents, query 4 was used to extract which compounds have the most connections to content entries and query 5 to count the unique food connections. The results of both queries are depicted in table 3 and table 4. Overall, the compounds contained in both tables are very similar and represent mainly essential elements and vitamins. This is not surprising as their contents in foods are generally well characterized in national nutritional databases.

**Table 3:** Compounds with highest associated contents count.

| Nr | Compound | Content Count |
| --- | --- | --- |
| 1 | retinol | 8184 |
| 2 | nicotinic acid | 7914 |
| 3 | sodium | 5757 |
| 4 | potassium | 5740 |
| 5 | iron | 5473 |
| 6 | phosphorus | 5180 |
| 7 | zinc | 5119 |
| 8 | riboflavin | 5016 |
| 9 | folic acid | 4878 |
| 10 | calcium | 4873 |
| 11 | copper | 4640 |
| 12 | pyridoxine | 4613 |
| 13 | pantothenic acid | 4567 |
| 14 | thiamine | 4555 |
| 15 | manganese | 4305 |

**Table 4:** Compounds with highest associated food counts.

| Nr | Compound | Food Count |
| --- | --- | --- |
| 1 | iron | 752 |
| 2 | calcium | 745 |
| 3 | phosphorus | 735 |
| 4 | sodium | 734 |
| 5 | potassium | 733 |
| 6 | riboflavin | 722 |
| 7 | copper | 719 |
| 8 | nicotinic acid | 717 |
| 9 | zinc | 715 |
| 10 | thiamine | 698 |
| 11 | manganese | 682 |
| 12 | pyridoxine | 673 |
| 13 | folic acid | 650 |
| 14 | selenium | 641 |
| 15 | retinol | 623 |

Both tables contain the elements sodium, potassium and phosphorus. These are important nutrients that patients on hemodialysis are advised to avoid overconsuming in their diet. The knowledge graph can be queried for foods that are very high and low in these minerals. For example, query 6 was used to extract the average potassium content in foods. The top 15 foods containing the most amount of potassium are depicted in table S16. In the first spot is kombu, which is a kelp belonging to the Laminariaceae family and a popular food in East Asia. Species from this family are known to contain high levels of many minerals, especially potassium [52, 53].

On the second spot in table S16 of foods with the most amount of potassium is the tomato. Fresh tomatoes are known to contain high amounts of potassium, with reported values ranging up to 1783 mg/100g [54]. However, the average value reported in table S16 is even higher with 3839.5 mg/100g and a large standard deviation of 1216.9 mg/100g. This equates to a potassium content of 3.8 % in tomato, which is doubtfully high. One advantage of the chosen graph structure is that each content entry also has a reference associated with it, which can be used to further elucidate this issue. Analysis of the information sources shows that the average value of 3839.5 mg/100g for the node “garden tomato (var.)” originates from two entries in the Dr. Duke’s Phytochemical and Ethnobotanical Database (Duke’s) [55], which is provided by the U.S. Department of Agriculture (USDA). However, due to the data contained in FooDB there is another node in the knowledge graph named “garden tomato”, which is connected to content measurements from the DTU and USDA‘s FoodData Central [56] which average around 796.5 mg/100g and 472.5 mg/100g respectively, around one fifth of the value from Duke’s.

The exact reason for the discrepancy between these measurements is speculative. While the cultivar type of the tomato might play a crucial role, potassium fertilization aimed at improving quality and yield could also lead to higher potassium content [57, 58]. However, most measured values mentioned in literature are closer to the FoodData content than to Duke’s. Interestingly, both Duke’s and FoodData Central are hosted by the USDA, yet they display significantly different values. This discrepancy underscores the importance of the knowledge graph in easily tracing statements back to their sources for comparison. It also highlights the considerable variability in measurement data, revealing that even for common compounds, information can be inconsistent. This demonstrates the need for continued efforts to enhance the accuracy and consistency of nutrient information.

Looking at top 15 foods with the lowest amount of potassium yields mostly processed foods and alcoholic beverages such as vodka, energy drink and marshmallow. These are not the types of food recommended to patients for a healthy diet. In this case the FoodOn ontology and RDF inference from GraphDB prove useful. By specifying that the returned foods should be of the type vegetables and vegetable products (FOODON_03540358) the adjusted query returns the vegetables with the lowest amounts of potassium, shown in table S17. These kinds of queries can serve as inspiration for healthy meal ingredients and, with future expansions of the knowledge graph to include recipes, enable recommendations for low-potassium dishes. Research indicates that maintaining adherence to optimal diets is still problematic [59–61], so providing a tool that simplifies meal planning would be very beneficial.

### 3.2 Exploration of the connection between compounds and conditions

Another crucial connection in the knowledge graph is through the measurement nodes that link compounds to conditions (see exemplary figure 4). Each measurement entry is associated with a compound and a condition as well as the sample type in which the compound was measured. Properties of the measurement nodes include the measured amount of the compound, the unit, and the reference, typically a PubMed ID for the publication. If available, additional information such as cohort or sex of the measured subjects is provided.

**Figure 4:**
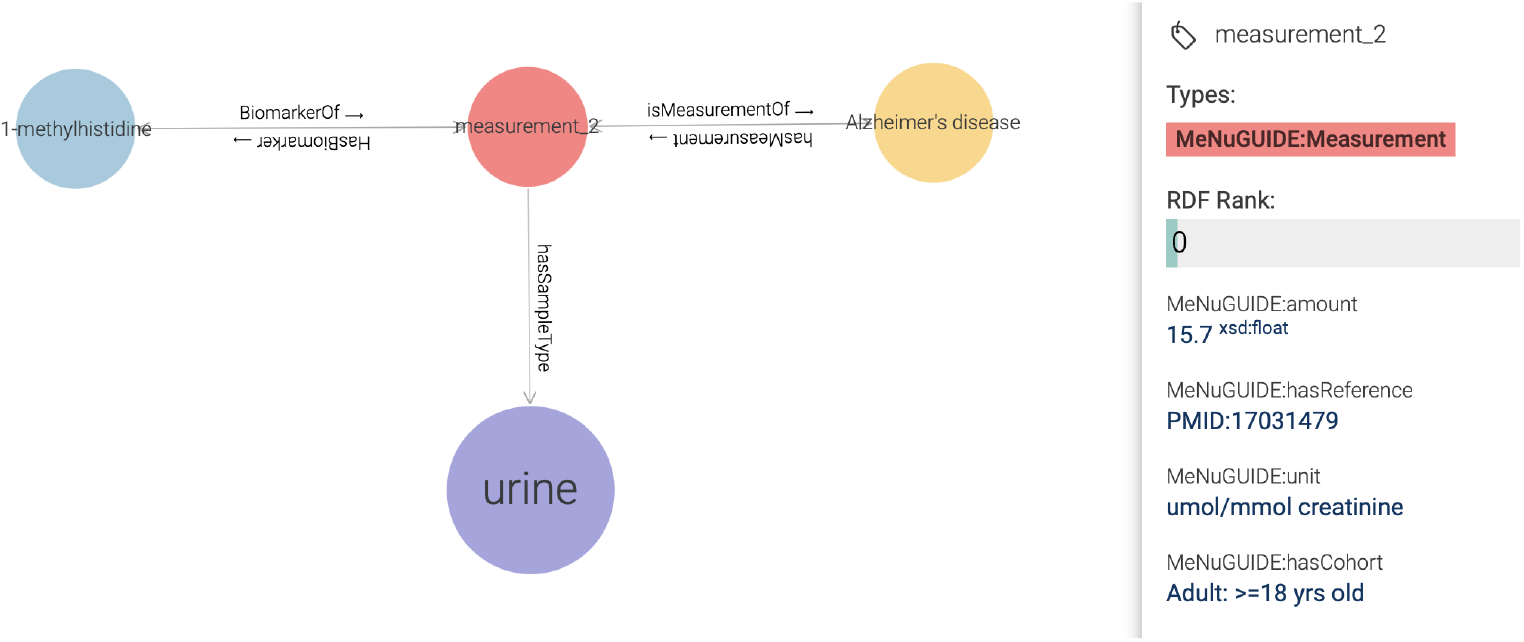
Example of a measurement node in MeNu GUIDE knowledge graph connecting compound and conditions.

Query 8 was used to extract which metabolites have the most connections to measurements, see table 5. Additionally, the number of connections to conditions via those measurement nodes was collected and is shown in table 6. Eleven of the 15 compounds found in table 5 also appear in table 6, which is much more consistent than the content associations. To further explore the connection between compounds and diseases in the knowledge graph, query 7 was used to find all conditions linked to a specific biomarker. Looking at l-lactic acid for example, the compound description already provides a first hint at the diseases it is involved in. The description states that “lactate measurement in critically ill patients has been traditionally used to stratify patients with poor outcomes”, which is likely a reason for why it is associated with so many measurements. Looking at the concentrations in different sample types, the knowledge graph contains measurements for urine, blood and cerebrospinal fluid. Many of the diseases found to be associated with l-lactic acid are inborn-errors of metabolism, such as Leigh disease, fructose-1,6-bisphosphatase deficiency and pyruvate decarboxylase deficiency, which the description already mentions. However, the query provides information on other linked diseases not mentioned in the description such as type 2 diabetes mellitus and epilepsy. At present, the reliance on MarkerDB results in relatively limited measurement connections that focus solely on biomarkers. However, literature mining could be utilized to broaden the scope and introduce additional causal relationships.

**Table 5:** Compounds with most associated measurements.

| Nr | Compound | Measurement Count |
| --- | --- | --- |
| 1 | l-lactic acid | 47 |
| 2 | uric acid | 33 |
| 3 | glycine | 28 |
| 4 | 3-hydroxybutyric acid | 25 |
| 5 | l-lysine | 24 |
| 6 | homovanillic acid | 21 |
| 7 | l-arginine | 21 |
| 8 | l-histidine | 20 |
| 9 | pipecolic acid | 20 |
| 10 | l-serine | 19 |
| 11 | cortisol | 19 |
| 12 | homocysteine | 18 |
| 13 | pyruvic acid | 17 |
| 14 | creatinine | 17 |
| 15 | hypoxanthine | 17 |

**Table 6:** Compounds with most associated conditions.

| Nr | Compound | Condition Count |
| --- | --- | --- |
| 1 | l-lactic acid | 20 |
| 2 | uric acid | 13 |
| 3 | l-arginine | 13 |
| 4 | glycine | 12 |
| 5 | homovanillic acid | 11 |
| 6 | 3-hydroxybutyric acid | 11 |
| 7 | homocysteine | 11 |
| 8 | pyruvic acid | 10 |
| 9 | xanthine | 10 |
| 10 | bilirubin | 10 |
| 11 | l-serine | 9 |
| 12 | l-lysine | 9 |
| 13 | l-alanine | 9 |
| 14 | creatinine | 9 |
| 15 | dopamine | 9 |

To complete the analysis of the knowledge graph connectivity, query 9 was used to extract the conditions that are connected to the most amount of measurements, seen in table 7. For comparison purposes, the conditions with the highest number of biomarkers associated via these measurement nodes were also extracted and are displayed in table 8. The only difference between these two results is that the condition maple syrup urine disease from table 7 is replaced with pre-eclampsia in table 8.

**Table 7:** Conditions with most associated measurements.

| Nr | Compound | Measurement Count |
| --- | --- | --- |
| 1 | normal | 1623 |
| 2 | obesity | 252 |
| 3 | Alzheimer's disease | 82 |
| 4 | eosinophilic esophagitis | 79 |
| 5 | uremia | 48 |
| 6 | chronic kidney disease | 28 |
| 7 | type 2 diabetes mellitus | 25 |
| 8 | Lewy body dementia | 21 |
| 9 | meningitis | 20 |
| 10 | schizophrenia | 19 |
| 11 | Cirrhosis | 19 |
| 12 | multiple sclerosis | 18 |
| 13 | Parkinson's disease | 18 |
| 14 | maple syrup urine disease | 18 |
| 15 | celiac disease | 17 |

**Table 8:**
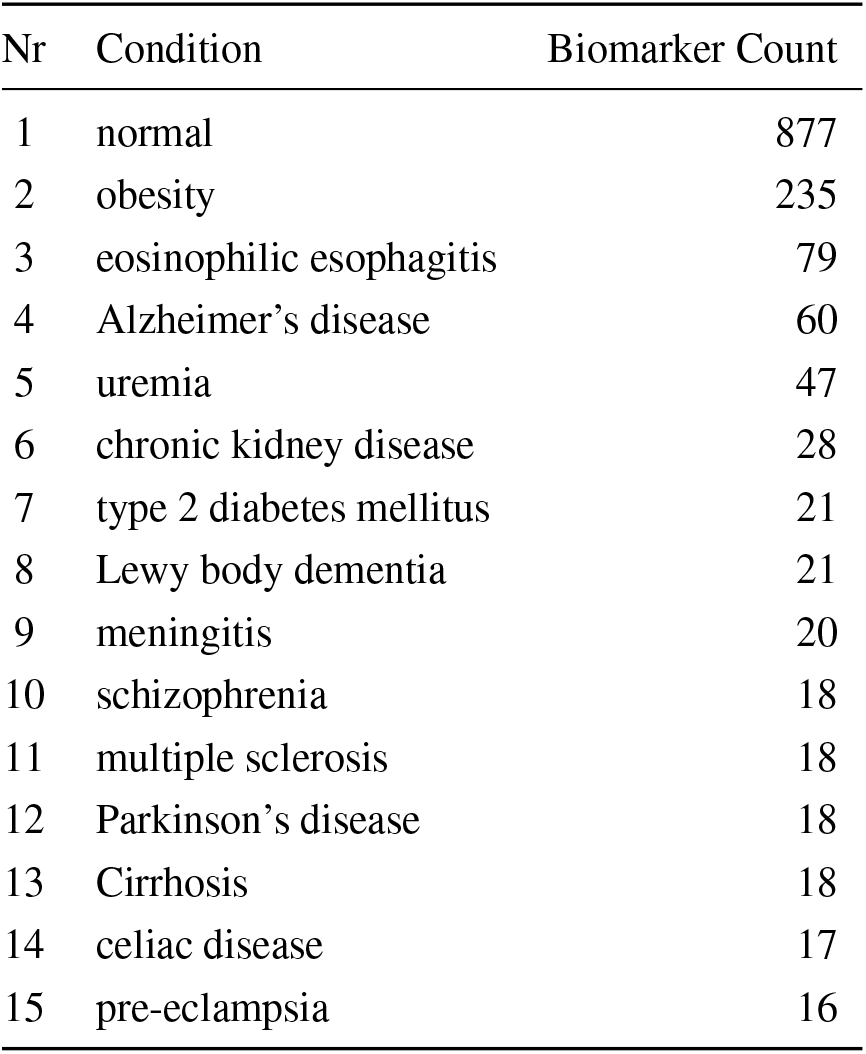
Conditions with most associated biomarkers.

| Nr | Condition | Biomarker Count |
| --- | --- | --- |
| 1 | normal | 877 |
| 2 | obesity | 235 |
| 3 | eosinophilic esophagitis | 79 |
| 4 | Alzheimer's disease | 60 |
| 5 | uremia | 47 |
| 6 | chronic kidney disease | 28 |
| 7 | type 2 diabetes mellitus | 21 |
| 8 | Lewy body dementia | 21 |
| 9 | meningitis | 20 |
| 10 | schizophrenia | 18 |
| 11 | multiple sclerosis | 18 |
| 12 | Parkinson's disease | 18 |
| 13 | Cirrhosis | 18 |
| 14 | celiac disease | 17 |
| 15 | pre-eclampsia | 16 |

Table 7 contains two diseases with notable diet associations: celiac disease and maple syrup urine disease. In the case of celiac disease, the critical compound, gluten, is unfortunately absent from the knowledge graph, as it is not part of the compounds extracted from the databases. Thus is cannot be connected to any food or condition. This is another clear knowledge gap that could be filled through text mining of relevant publications. However, for maple syrup urine disease, leucine and isoleucine were identified as biomarkers. Individuals with this disease cannot properly catabolize the branched-chain amino acids leucine, isoleucine, and valine due to a genetic mutation.

This association now can serve as the connection to enable the main feature of the knowledge graph, which is the extraction of foods that contain the compounds connected to a specific disease. The knowledge graph facilitates the direct creation of a connection between the condition, its associated biomarkers, and the foods that contain them, as demonstrated by query 10. This relationship can be visualized using the “CONSTRUCT” keyword in the SPARQL query 11, with figure 5 showing an exemplary visualization for the top three foods with a link display limit of 50.

**Figure 5:**
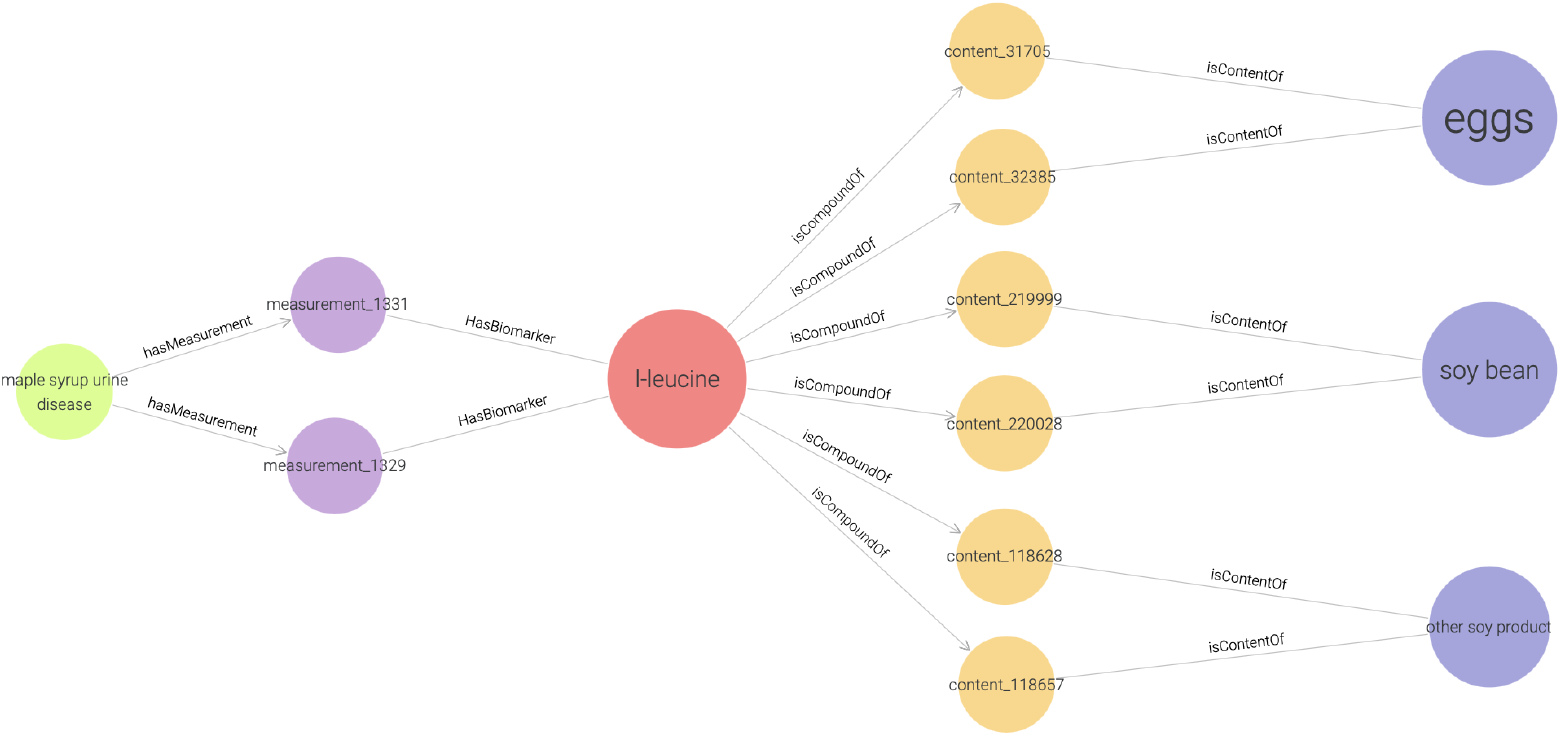
Example of condition-compound-food connection in the MeNu GUIDE knowledge graph showing the top three foods in the knowledge graph containing the most amount of l-leucine, which is a biomarker for maple syrup urine disease.

Each of the top three foods (eggs, soy bean and other soy product) is connected to the l-leucine node via two content nodes that contain reference measurements from the USDA database. The compound node of l-leucine is then connected to maple syrup urine disease via two measurement nodes, measurement_1331 and measurement_1329. The node properties (figure S1 A) reveal that the source of measurement_1331 is a publication from 2002 that measured an average blood concentration of 655 *μM* l-leucine levels in newborns with maple syrup urine disease [62]. Analogously, the properties of the measurement_1329 node (figure S1 B) point to a publication from 1980 where a l-leucine level of 892 *μM* was measured in the cerebrospinal fluid of a two year old child [63].

This example demonstrates the capability of the knowledge graph to reveal critical connections and present them in a visually engaging and easily navigable format. Additionally, by providing the source for each connection within the graph researchers are able to quickly locate publications supporting specific data points or relationships. This not only enables the fast extraction of relevant literature on a certain research question, but also easy verification of the contained data. Most notably though, a single query was utilized to identify compounds associated with a specific disease and to pinpoint which foods contain these compounds. Achieving the same result without the knowledge graph would involve laborious manual searches across multiple databases. This functionality underscores a significant advantage of the knowledge graph, made possible through the successful integration of diverse data sources in an ontology-based framework.

## 4 Conclusion and Outlook

The MeNu GUIDE knowledge graph serves as a proof of concept, demonstrating the feasibility of linking food and disease at a metabolite-specific level to extract relevant information. For instance, the graph structure facilitates the identification of compounds shared by different foods, allowing for the investigation of metabolic changes that occur during food processing, such as cheese-making. Additionally, the knowledge graph enables users to search for foods with particularly high or low levels of specific compounds, aiding in dietary decisions to either boost essential nutrients or avoid excessive consumption of unfavorable substances. Most notably, a single query can reveal compounds associated with a particular disease and locate foods that contain them. This functionality underscores a significant benefit of the knowledge graph, made possible by the successful integration of various data sources.

The MeNu GUIDE knowledge graph establishes a foundation by offering a standardized, ontology-based framework, which can easily be expanded. The current implementation provides a solid starting point for developing increasingly complex models. The analyses revealed that relying solely on database information often results in significant knowledge gaps. For instance, crucial details such as the role of gluten in celiac disease are missing. Therefore, the primary task moving forward will be to enhance the knowledge graph with additional information. This can include data from other databases as well as insights from scientific literature, which harbors a wealth of untapped knowledge. However, accessing this valuable information is challenging due to its fragmented nature, and manually extracting relevant details would be exceedingly time-consuming. Modern technologies, such as neural networks and machine learning algorithms, present a promising solution to this problem [64–67]. Additionally, incorporating recipes into the database would facilitate the development of a recommendation algorithm tailored to specific nutritional needs. Including information on bioavailability during this process would also be highly beneficial.

The second major objective going forward is validation. Once additional content is integrated, the knowledge graph should be made available as a web application for the scientific community. This will allow researchers and experts to identify and report errors for correction. Given the vast scale of the knowledge graph, containing millions of edges even before incorporating data from literature, it is impractical for a single individual alone to validate the content. Therefore, a collaborative approach is essential. Similar to Wikipedia, experts in relevant fields should have the ability to report issues and make corrections, leveraging the collective intelligence of the community.

By expanding the content, facilitating validation, and providing an intuitive framework for exploration, the knowledge graph has the potential to become an invaluable tool for researchers. It allows to explore the currently available knowledge on a topic and discover gaps that might serve as interesting areas for further investigation. Empowering researchers with a comprehensive knowledge graph has the potential to catalyze breakthroughs in understanding the intricate connections between diet and health. As nutrition research evolves, there is a growing need for a tool that aggregates significant findings and makes them readily accessible and searchable. This pilot project has demonstrated that an ontology-based knowledge graph is well-suited to meet this need.

## Data Availability Statement

All data used for this project is publicly available. The Turtle file containing the complete knowledge graph can be downloaded from figshare via https://doi.org/10.6084/m9.figshare.27216054.v1.

## Code Availability Statement

The MeNu GUIDE source code can be accessed on GitHub via https://github.com/LipiTUM/MeNuGUIDE.

## Author Contributions

JKP conceived the initial idea, supervised the project and secured the funding. VW and JKP planned and conceptualized the work. VW extracted the database information, merged the ontologies and ran all queries. VW and JKP wrote the manuscript. All authors read, reviewed, and accepted the manuscript in its final form.

## Acknowledgments

This project was funded by the Bavarian State Ministry of Science and the Arts in the framework of the Bavarian Research Institute for Digital Transformation (bidt; JKP, VW: Junior Research Group LipiTUM).

## List of Abbreviations

CDNO: Compositional Dietary Nutrition Ontology
ChEBI: Chemical Entities of Biological Interest
CHIRO: ChEBI Integrated Role Ontology
DO: Human Disease Ontology
DrOn: The Drug Ontology
DTU: Technical University of Denmark
Duke’s: Dr. Duke’s Phytochemical and Ethnobotanical Database
FIDEO: Food Interactions with Drugs Evidence Ontology
FOBI: Food-Biomarker Ontology
FooDB: Food Database
FoodOn: Food Ontology
GO: Gene Ontology
HMDB: Human Metabolome Database
IBD: Inflammatory Bowel Disease
IBS: Irritable Bowel Syndrome
InChI: International Chemical Identifier
IRI: Internationalized Resource Identifier
KEGG: Kyoto Encyclopedia of Genes and Genomes
MCDB: Milk Composition Database
RDF: Resource Description Framework
SBO: Systems Biology Ontology
SPARQL: SPARQL Protocol and RDF Query Language
USDA: U.S. Department of Agriculture
VMH: Virtual Metabolic Human

