## Supplements for "MeNu GUIDE - a metabolite nutrition graph to uncover interactions with disease etiology"

### Supplement

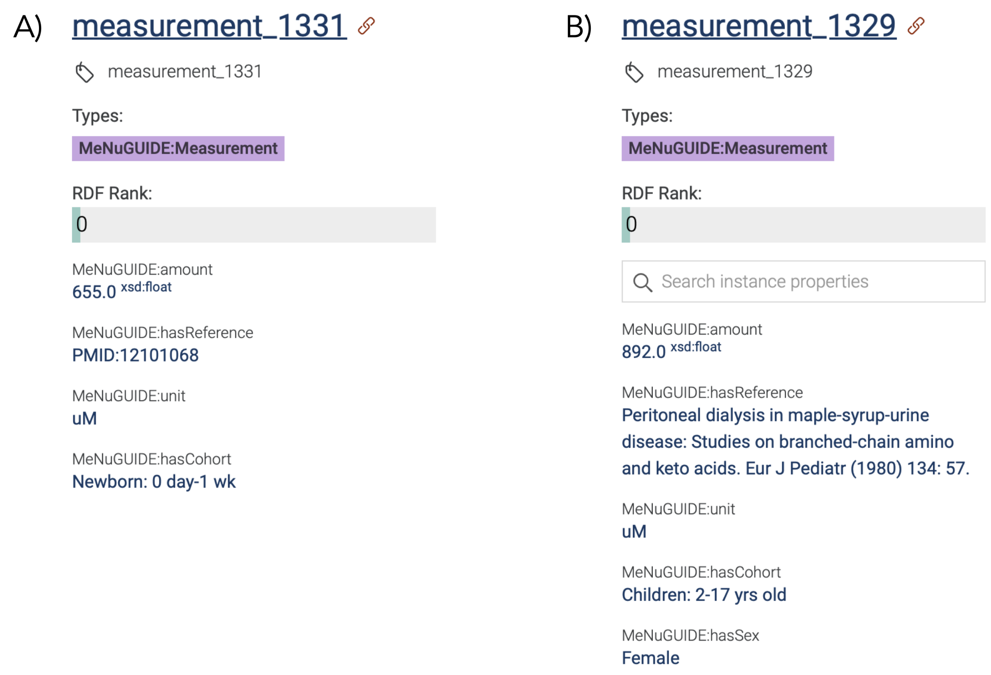


***Figure S1:*** *Properties associated with the measurement nodes A) measurement_1331 and B) measurement_1329.*

***Table S9:*** *Properties of compounds and their corresponding triple predicates.*

| Object | Predicate |
|---|---|
| Compound Name | http://www.w3.org/2000/01/rdf-schema#label |
| Description | http://purl.obolibrary.org/obo/IAO_0000115 |
| Chemical Formula | http://purl.obolibrary.org/obo/chebi/formula |
| Monoisotopic Mass | http://purl.obolibrary.org/obo/chebi/monoisotopicmass |
| SMILES | http://purl.obolibrary.org/obo/chebi/smiles |
| InChI | http://purl.obolibrary.org/obo/chebi/inchi |
| InChIKey | http://purl.obolibrary.org/obo/chebi/inchikey |
| IUPAC Name | http://www.geneontology.org/formats/oboInOwl#hasExactSynonym |
| CAS Number | http://www.geneontology.org/formats/oboInOwl#hasDbXref |
| HMDB ID | http://www.geneontology.org/formats/oboInOwl#hasDbXref |
| FooDB ID | http://www.geneontology.org/formats/oboInOwl#hasDbXref |
| KEGG ID | http://www.geneontology.org/formats/oboInOwl#hasDbXref |
| VMH ID | http://www.geneontology.org/formats/oboInOwl#hasDbXref |
| MarkerDB ID | http://www.geneontology.org/formats/oboInOwl#hasDbXref |
| Exposome Explorer ID | http://www.geneontology.org/formats/oboInOwl#hasDbXref |
| ChEBI Class IRI | http://www.w3.org/1999/02/22-rdf-syntax-ns#type |
| MeNu GUIDE Compound | http://www.w3.org/1999/02/22-rdf-syntax-ns#type |
| Reaction IRI (input of) | http://purl.obolibrary.org/obo/RO_0002352 |
| Reaction IRI (output of) | http://purl.obolibrary.org/obo/RO_0002353 |
| Content IRI | http://MeNuGUIDE.local/isCompoundOf |
| Measurement IRI (BiomarkerOf) | http://purl.obolibrary.org/obo/FOBI_00422 |

***Table S10:*** *Properties of foods and their corresponding triple predicates.*

| Object | Predicate |
|---|---|
| Food Name | http://www.w3.org/2000/01/rdf-schema#label |
| Scientific Name | http://www.geneontology.org/formats/oboInOwl#hasSynonym |
| Description | http://purl.obolibrary.org/obo/IAO_0000115 |
| FooDB ID | http://www.geneontology.org/formats/oboInOwl#hasDbXref |
| FoodON IRI | http://www.w3.org/1999/02/22-rdf-syntax-ns#type |
| NCBI IRI | http://www.w3.org/1999/02/22-rdf-syntax-ns#type |
| MeNu GUIDE Food | http://www.w3.org/1999/02/22-rdf-syntax-ns#type |
| Content IRI | http://MeNuGUIDE.local/hasContent |

***Table S11:*** *Properties of content measurements and their corresponding triple predicates.*

| Object | Predicate |
|---|---|
| Content ID | http://www.w3.org/2000/01/rdf-schema#label |
| MeNu GUIDE Content | http://www.w3.org/1999/02/22-rdf-syntax-ns#type |
| Food IRI | http://MeNuGUIDE.local/isContentOf |
| Compound IRI | http://MeNuGUIDE.local/hasCompound |
| Quantity | http://MeNuGUIDE.local/amount |
| Minimum quantity | http://MeNuGUIDE.local/minimumAmount |
| Maximum quantity | http://MeNuGUIDE.local/maximumAmount |
| Unit | http://MeNuGUIDE.local/unit |
| Reference | http://MeNuGUIDE.local/hasReference |

***Table S12:*** *Properties of reactions and their corresponding triple predicates.*

| Object | Predicate |
|---|---|
| Reaction Abbreviation | http://www.w3.org/2000/01/rdf-schema#label |
| Description | http://purl.obolibrary.org/obo/IAO_0000115 |
| KEGG ID | http://www.geneontology.org/formats/oboInOwl#hasDbXref |
| VMH ID | http://www.geneontology.org/formats/oboInOwl#hasDbXref |
| MeNu GUIDE Reaction | http://www.w3.org/1999/02/22-rdf-syntax-ns#type |
| Compound IRI (has input) | http://purl.obolibrary.org/obo/RO_0002233 |
| Compound IRI (has output) | http://purl.obolibrary.org/obo/RO_0002234 |
| Enzyme IRI (has participant) | http://purl.obolibrary.org/obo/RO_0000057 |

***Table S13:*** *Properties of enzymes and their corresponding triple predicates.*

| Object | Predicate |
|---|---|
| EC Number | http://www.w3.org/2000/01/rdf-schema#label |
| MeNu GUIDE Enzyme | http://www.w3.org/1999/02/22-rdf-syntax-ns#type |
| Reaction IRI (participates in) | http://purl.obolibrary.org/obo/RO_0000056 |
| GO Term IRI (has function) | http://purl.obolibrary.org/obo/RO_0000085 |

***Table S14:*** *Properties of genes and their corresponding triple predicates.*

| Object | Predicate |
|---|---|
| Gene Symbol | http://www.w3.org/2000/01/rdf-schema#label |
| MeNu GUIDE Gene | http://www.w3.org/1999/02/22-rdf-syntax-ns#type |
| Reaction IRI (involved in) | http://purl.obolibrary.org/obo/RO_0002331 |

***Table S15:*** *Properties of biomarker measurements and their corresponding triple predicates.*

| Object | Predicate |
|---|---|
| Measurement ID | http://www.w3.org/2000/01/rdf-schema#label |
| MeNu GUIDE Measurement | http://www.w3.org/1999/02/22-rdf-syntax-ns#type |
| Condition IRI | http://MeNuGUIDE.local/isMeasurementOf |
| Compound IRI (hasBiomarker) | http://purl.obolibrary.org/obo/FOBI_00423 |
| Quantity | http://MeNuGUIDE.local/amount |
| Unit | http://MeNuGUIDE.local/unit |
| Cohort | http://MeNuGUIDE.local/hasCohort |
| Sex | http://MeNuGUIDE.local/hasSex |
| Sample Type | http://MeNuGUIDE.local/hasSampleType |
| Reference | http://MeNuGUIDE.local/hasReference |

***Table S16:*** *Foods with highest amounts of potassium in mg/100g.*

| Nr | Food | Content | SD |
|---|---|---|---|
| 1 | kombu | 7500.0 | |
| 2 | garden tomato (var.) | 3839.5 | 1216.9 |
| 3 | scarlet bean | 3580.7 | |
| 4 | cornmint | 3510.0 | |
| 5 | chervil | 3422.3 | 2448.7 |
| 6 | leavening agent | 3344.0 | 5526.2 |
| 7 | tarragon | 3078.5 | 82.7 |
| 8 | mugwort | 2910.0 | 1004.1 |
| 9 | endive | 2629.9 | 3275.1 |
| 10 | wild carrot | 2468.0 | |
| 11 | turmeric | 2416.0 | 154.1 |
| 12 | lambsquarters | 2358.0 | 3574.5 |
| 13 | parsley | 2275.5 | 2251.2 |
| 14 | cabbage | 2243.4 | |
| 15 | sweet basil | 2228.7 | 1767.5 |

***Table S17:*** *Vegetables with lowest amounts of potassium in mg/100g.*

| Nr | Compound | Content | SD |
|---|---|---|---|
| 1 | wakame | 50.0 | |
| 2 | horned melon | 123.0 | |
| 3 | green bean | 182.5 | 75.1 |
| 4 | black cabbage | 197.3 | 66.8 |
| 5 | okra | 208.3 | 70.6 |
| 6 | welsh onion | 212.0 | |
| 7 | rice | 212.1 | 347.8 |
| 8 | american pokeweed | 213.0 | 41.0 |
| 9 | cauliflower | 216.5 | 83.9 |
| 10 | corn | 229.5 | 202.3 |
| 11 | savoy cabbage | 230.0 | |
| 12 | leek | 236.0 | |
| 13 | butternut squash | 245.3 | 94.2 |
| 14 | ostrich fern | 249.5 | 170.4 |
| 15 | broccoli | 274.8 | 192.7 |

```sparql
PREFIX rdfs: <http://www.w3.org/2000/01/rdf-schema#>
PREFIX rdf: <http://www.w3.org/1999/02/22-rdf-syntax-ns#>
PREFIX MeNuGUIDE: <http://MeNuGUIDE.local/>

SELECT ?food (COUNT(?object) as ?count) ?label
WHERE {
    ?food rdf:type MeNuGUIDE:Food .
    ?food rdfs:label ?label .
    ?food MeNuGUIDE:hasContent ?object .
}
GROUP BY ?food ?label
ORDER BY DESC(?count)
LIMIT 25
```

***Listing 1:*** *Top 25 Foods by Content Measurements*

```sparql
PREFIX rdfs: <http://www.w3.org/2000/01/rdf-schema#>
PREFIX rdf: <http://www.w3.org/1999/02/22-rdf-syntax-ns#>
PREFIX MeNuGUIDE: <http://MeNuGUIDE.local/>
PREFIX agg: <http://jena.apache.org/ARQ/function/aggregate#>

SELECT ?compoundName (AVG(?amount) as ?average_amount) ?unit (agg:stdev(?amount) as ?
    stddev)
WHERE {
    ?food rdf:type MeNuGUIDE:Food .
    ?food rdfs:label "cattle (beef, veal)" .
    ?food MeNuGUIDE:hasContent ?content .
    ?content MeNuGUIDE:hasCompound ?compound .
    ?compound rdfs:label ?compoundName .
    ?content MeNuGUIDE:amount ?amount .
    ?content MeNuGUIDE:unit ?unit .
}
GROUP BY ?compoundName ?unit
ORDER BY DESC(?average_amount)
```

***Listing 2:*** *Average compound content in food node cattle*

```sparql
PREFIX rdfs: <http://www.w3.org/2000/01/rdf-schema#>
PREFIX rdf: <http://www.w3.org/1999/02/22-rdf-syntax-ns#>
PREFIX MeNuGUIDE: <http://MeNuGUIDE.local/>

SELECT ?compound ?compoundName (AVG(?content1amount) as ?averageContent1Amount) (AVG
    (?content2amount)  as ?averageContent2Amount)
WHERE {
    ?compound rdf:type MeNuGUIDE:Compound .
    ?compound rdfs:label ?compoundName .

    ?food1 rdfs:label "milk (cow)" .
    ?food2 rdfs:label "cheese" .

    ?food1 MeNuGUIDE:hasContent ?content1 .
    ?content1 MeNuGUIDE:hasCompound ?compound .
    ?content1 MeNuGUIDE:amount ?content1amount .
    ?content1 MeNuGUIDE:unit "mg/100g" .

    ?food2 MeNuGUIDE:hasContent ?content2 .
    ?content2 MeNuGUIDE:hasCompound ?compound .
    ?content2 MeNuGUIDE:amount ?content2amount .
    ?content2 MeNuGUIDE:unit "mg/100g" .

}
GROUP BY ?compound ?compoundName
LIMIT 25
```

***Listing 3:*** *Comparison of Compound Contents between Cow Milk and Cheese*

```
PREFIX rdfs: <http://www.w3.org/2000/01/rdf-schema#>
PREFIX rdf: <http://www.w3.org/1999/02/22-rdf-syntax-ns#>
PREFIX MeNuGUIDE: <http://MeNuGUIDE.local/>

SELECT ?compound (COUNT(?object) as ?count) ?label
WHERE {
    ?compound rdf:type MeNuGUIDE:Compound .
    ?compound rdfs:label ?label .
    ?compound MeNuGUIDE:isCompoundOf ?object .
}
GROUP BY ?compound ?label
ORDER BY DESC(?count)
LIMIT 25
```

***Listing 4:*** *Top 25 Compounds by Content Measurements*

```
PREFIX rdfs: <http://www.w3.org/2000/01/rdf-schema#>
PREFIX rdf: <http://www.w3.org/1999/02/22-rdf-syntax-ns#>
PREFIX MeNuGUIDE: <http://MeNuGUIDE.local/>

SELECT ?compoundName (COUNT(DISTINCT ?food) as ?foodCount)
WHERE {
    ?compound rdf:type MeNuGUIDE:Compound .
    ?compound rdfs:label ?compoundName .
    ?compound MeNuGUIDE:isCompoundOf ?content .
    ?content MeNuGUIDE:isContentOf ?food .
    ?food rdfs:label ?foodName .
}
GROUP BY ?compoundName
ORDER BY DESC(?foodCount)
LIMIT 15
```

***Listing 5:*** *Top 15 Compounds by Food Count*

```
PREFIX rdfs: <http://www.w3.org/2000/01/rdf-schema#>
PREFIX rdf: <http://www.w3.org/1999/02/22-rdf-syntax-ns#>
PREFIX MeNuGUIDE: <http://MeNuGUIDE.local/>

SELECT ?food ?foodName (AVG(?amount) as ?average)
WHERE {
    ?food rdf:type MeNuGUIDE:Food .
    ?food MeNuGUIDE:hasContent ?content .
    ?food rdfs:label ?foodName .
    ?content MeNuGUIDE:hasCompound ?compound .
    ?content MeNuGUIDE:amount ?amount .
    ?content MeNuGUIDE:unit "mg/100g" .
    ?compound rdf:type MeNuGUIDE:Compound .
    ?compound rdfs:label "potassium" .
}
GROUP BY ?food ?foodName
ORDER BY DESC(?average)
```

***Listing 6:*** *Average Amount of Potassium in Foods in mg/100g*

```
PREFIX obo: <http://purl.obolibrary.org/obo/>
PREFIX rdfs: <http://www.w3.org/2000/01/rdf-schema#>
PREFIX rdf: <http://www.w3.org/1999/02/22-rdf-syntax-ns#>
PREFIX MeNuGUIDE: <http://MeNuGUIDE.local/>

SELECT ?conditionName (AVG(?content) as ?meanContent) ?unit ?sampleTypeName
WHERE {
    ?compound rdf:type MeNuGUIDE:Compound .
    ?compound rdfs:label "l-lactic acid" .
    ?compound obo:IAO_0000115 ?description .
    ?compound obo:FOBI_00422 ?measurement .
    ?measurement MeNuGUIDE:isMeasurementOf ?condition .
    ?measurement MeNuGUIDE:amount ?content .
    ?measurement MeNuGUIDE:unit ?unit .
    ?measurement MeNuGUIDE:hasSampleType ?sampleType .
    ?sampleType rdfs:label ?sampleTypeName .
    ?condition rdfs:label ?conditionName .
}
GROUP BY ?conditionName ?unit ?sampleTypeName
ORDER BY ?meanContent
```
***Listing 7:*** *Average concentration of l-lactic acid in different sample types associated with various conditions*

```
PREFIX obo: <http://purl.obolibrary.org/obo/>
PREFIX rdfs: <http://www.w3.org/2000/01/rdf-schema#>
PREFIX rdf: <http://www.w3.org/1999/02/22-rdf-syntax-ns#>
PREFIX MeNuGUIDE: <http://MeNuGUIDE.local/>

SELECT ?compoundName (COUNT(?measurement) as ?count)
WHERE {
    ?compound rdf:type MeNuGUIDE:Compound .
    ?compound rdfs:label ?compoundName .
    ?compound obo:FOBI_00422 ?measurement .
}
GROUP BY ?compoundName
ORDER BY DESC(?count)
LIMIT 15
```
***Listing 8:*** *Top 25 Compounds by Biomarker Measurements*

```
PREFIX rdfs: <http://www.w3.org/2000/01/rdf-schema#>
PREFIX rdf: <http://www.w3.org/1999/02/22-rdf-syntax-ns#>
PREFIX MeNuGUIDE: <http://MeNuGUIDE.local/>

SELECT ?condition (COUNT(?object) as ?count) ?label
WHERE {
    ?condition rdf:type MeNuGUIDE:Condition .
    ?condition rdfs:label ?label .
    ?condition MeNuGUIDE:hasMeasurement ?object .
}
GROUP BY ?condition ?label
ORDER BY DESC(?count)
LIMIT 25
```
***Listing 9:*** *Top 25 Conditions by Biomarker Measurements*

```
PREFIX obo: <http://purl.obolibrary.org/obo/>
PREFIX rdfs: <http://www.w3.org/2000/01/rdf-schema#>
PREFIX rdf: <http://www.w3.org/1999/02/22-rdf-syntax-ns#>
PREFIX MeNuGUIDE: <http://MeNuGUIDE.local/>
PREFIX agg: <http://jena.apache.org/ARQ/function/aggregate#>

SELECT ?compoundName ?foodName (AVG(?amount) as ?averageAmount) (agg:stdev(?amount)
    as ?stddev)
WHERE {
    ?condition rdf:type MeNuGUIDE:Condition .
    ?condition rdfs:label "maple syrup urine disease"@en .
    ?condition MeNuGUIDE:hasMeasurement ?measurement .

    ?measurement obo:FOBI_00423 ?compound .

    ?compound rdfs:label ?compoundName .
    ?compound MeNuGUIDE:isCompoundOf ?content .

    ?content MeNuGUIDE:isContentOf ?food .
    ?content MeNuGUIDE:amount ?amount .
    ?content MeNuGUIDE:unit "mg/100g" .

    ?food rdfs:label ?foodName .
}
GROUP BY ?compoundName ?foodName
ORDER BY DESC(?averageAmount)
```
***Listing 10:*** *Biomarkers associated with maple syrup urine disease and they content in foods*

```
PREFIX obo: <http://purl.obolibrary.org/obo/>
PREFIX rdfs: <http://www.w3.org/2000/01/rdf-schema#>
PREFIX rdf: <http://www.w3.org/1999/02/22-rdf-syntax-ns#>
PREFIX MeNuGUIDE: <http://MeNuGUIDE.local/>
PREFIX xsd: <http://www.w3.org/2001/XMLSchema#>

CONSTRUCT {
    ?compound a MeNuGUIDE:Compound ;
              rdfs:label "l-leucine";
              MeNuGUIDE:isCompoundOf ?content .

    ?food a MeNuGUIDE:Food ;
          rdfs:label ?foodName .

    ?condition a MeNuGUIDE:Condition ;
               rdfs:label "maple syrup urine disease"@en ;
               MeNuGUIDE:hasMeasurement ?measurement .

    ?measurement a MeNuGUIDE:Measurement ;
                 obo:FOBI_00423 ?compound .

    ?content a MeNuGUIDE:Content ;
         MeNuGUIDE:isContentOf ?food ;
             MeNuGUIDE:amount ?amount ;
         MeNuGUIDE:unit "mg/100g" ;
         MeNuGUIDE:hasReference ?reference .

} WHERE {
    ?condition rdf:type MeNuGUIDE:Condition .
    ?condition rdfs:label "maple syrup urine disease"@en .
    ?condition MeNuGUIDE:hasMeasurement ?measurement .

    ?measurement obo:FOBI_00423 ?compound .

    ?compound rdfs:label "l-leucine" .
    ?compound MeNuGUIDE:isCompoundOf ?content .

    ?content MeNuGUIDE:isContentOf ?food .
    ?content MeNuGUIDE:amount ?amount .
    ?content MeNuGUIDE:unit "mg/100g" .
    ?content MeNuGUIDE:hasReference ?reference .

    ?food rdfs:label ?foodName .

    FILTER (!CONTAINS(STR(?reference), "DTU"))
}
ORDER BY DESC(?amount)
```

***Listing 11:*** *Construct graph of biomarkers associated with maple syrup urine disease and they content in foods*
